# A metagenomic exploration of the bacterial community composition of two deep-sea *Pheronema carpenteri* sponge aggregations from the North Atlantic; insights into ecosystem services

**DOI:** 10.64898/2026.03.27.714666

**Authors:** Poppy J. Hesketh-Best, Matthew J. Koch, Nicola L. Foster, Philip J. Warburton, Mathew Upton, Kerry Howell

## Abstract

**Aims:** Sponge microbiomes have been extensively studied, in part due to their potential as sources of novel antimicrobials and other biologics, with most research focusing on Demosponges. Here, we investigate the Hexactinellid sponge *Pheronema carpenteri*, previously identified as a promising source of antibiotic-producing bacteria.

**Methods:** Using next-generation sequencing of bacterial 16S rRNA genes and a single sponge metagenome, we examined the composition of bacterial communities of *P. carpenteri* sponges recovered from the Porcupine Seabight, along with local water and sediment samples.

**Results:** Our results show that *P. carpenteri* harbours a microbiome abundant in Proteobacteria (47.1-59.4%) and Actinobacteria (11.5-27.5%), with consistent intra-aggregation similarities and structured intra-sponge communities.

A metagenomic analysis revealed the presence of several nitrogen cycling genes (*nirK, nosZ, nirS* homologues of proteobacterial origin), supporting a suggestion that these sponges may play a role in nitrogen cycling, while biosynthetic gene clusters (BGCs) were limited (4 complete clusters). Notably, bacterial community structures within *P. carpenteri* aggregations resemble those observed in both low and high microbial abundance (LMA/HMA) sponges.

**Conclusions:** Hexactinellids are traditionally considered LMA sponges, so identifying species that deviate from this dichotomy provides new insights into sponge microbiome ecology. Integrating Hexactinellids into both culture-dependent and culture-independent studies will advance our broader understanding of sponge-associated microbial diversity and could inform biodiscovery programmes in marine environments.

**Impact Statement:** Our findings support the suggestion that a combination of culture-based and molecular analyses is required to generate a comprehensive picture of the biosynthetic potential of *P. carpenteri* sponges. We also reveal insights into the ecosystem services that sponge microbiomes may contribute towards. These observations could facilitate a deeper understanding of the biotechnological and environmental value of key marine resources.

## Introduction

Sponges are key components of marine benthic ecosystems, providing essential functions such as nutrient cycling, habitat formation, fish refuge, and a source of novel chemical compounds (Bell 2008; Goeij *et al*. 2013; Maldonado 2016; Conway *et al*. 2025). In the deep sea, sponges can form dense aggregations of single species, such as *Pheronema carpenteri* (Howell *et al*. 2016), which are of conservation importance (ICES, 2016). The ecological roles of deep-sea sponges are closely linked to the complex associations that sponges form with microbial symbionts, with up to ∼71 bacterial phyla and multiple archaeal and eukaryotic lineages detected in deep-sea species (Thomas *et al*. 2016; Moitinho-Silva *et al*. 2017; Steinert *et al*. 2020, 2020).

Hexactinellida (glass sponges) are predominantly deep-sea organisms, occurring at depths >200 m (Leys *et al*. 2004). Dense aggregations of *P. carpenteri* have been recorded at up to 1.53 individuals m^−2^ at ∼1,200 m in the North-East Atlantic (Hughes and Gage 2004). Historically, Hexactinellida have been underrepresented in microbiome studies and considered low microbial abundance (LMA) species, in contrast to Demospongiae where a high (HMA) vs. low (LMA) microbial abundance dichotomy has been established as a key indicator of microbial composition (Weisz, Lindquist and Martens 2008; Giles *et al*. 2013; Erwin *et al*. 2015). Recent analyses indicate that Hexactinellida largely exhibit LMA-like microbiomes, although some lineages display greater complexity resembling HMA profiles, suggesting class-specific trends in symbiont assembly and functional specialization (Turon *et al*. 2019; Busch *et al*. 2022; Díez-Vives and Riesgo 2024). Despite their ecological importance, the microbiomes of Hexactinellida are still not well understood and show a more uneven microbial distribution compared to other LMA sponges (Tian *et al*. 2016; Bayer *et al*. 2020; Steinert *et al*. 2020). Notably, some LMA hexactinellid species display Shannon diversity levels comparable to those of HMA sponges (Busch *et al*. 2022).

Current knowledge of Hexactinellida microbiomes derives from culture-based studies (Mangano *et al*. 2008; Xin *et al*. 2011; Koch *et al*. 2021; Conway *et al*. 2025), 16S rRNA surveys (Thomas *et al*. 2016; Steinert *et al*. 2020), and metagenomic reconstructions (Tian *et al*. 2016; Bayer *et al*. 2020; Wei *et al*. 2023). Culture-based approaches have identified *P. carpenteri* as promising for natural product discovery (Koch *et al*. 2021; Hesketh-Best *et al*. 2023; Conway *et al*. 2025), but cultivation captures only a fraction of microbial diversity, leaving considerable scope for exploring the genomic potential. Phylogenetic surveys indicate that Hexactinellida microbial communities resemble surrounding seawater, dominated by Proteobacteria, Bacteroidetes, Chloroflexi, and ammonia-oxidizing Nitrosopumilaceae, with alpha diversity lower than Demospongiae (Bayer *et al*. 2020: 20; Steinert *et al*. 2020). Vertical gradients (depth) appear to influence microbial composition more than horizontal geographic variation (Sunagawa *et al*. 2015; Li *et al*. 2018), consistent with the similarity between Hexactinellida and surrounding water column communities (Thomas *et al*. 2016; Steinert *et al*. 2020).

Despite their ecological and biotechnological potential, deep-water Hexactinellida remain underexplored for natural product discovery. Culture-dependent studies have yielded bioactive strains from *Sericolophus hawaiicus* and *Rossella* spp., including *Dietzia* sp. with polyketide synthase (PKS) genes and *Pseudoalteromonas* sp. with high protease activity (Xin *et al*. 2011; Borchert *et al*. 2016). Here, we investigate the microbiome of *P. carpenteri*, testing for (i) differences between sponge-associated and environmental microbial communities, (ii) inter-site- and intra-site variation in sponge microbiota, and (iii) microbial functional potential using nitrogen cycling gene analyses.

## Materials and Methods

### Collection of sponges and environmental samples

*P. carpenteri* sponges were collected in August 2019 from the Porcupine Seabight, North-East Atlantic, during the SeaRovers 2019 research cruise on the RV *Celtic Explorer* (CE19-15-029). Using the ROV *Holland I*, sponges were sampled alongside sediment and seawater at depths of 1,103 and 1,208 m on the east and west sides of the Seabight (Figure 1, Supp.Table 1). At sites T7 and T52, three sponges per site, sediment samples, and 1.5 L of seawater (collected via Niskin bottles) were obtained. On return of the ROV to the ship, water samples were filtered through 0.22 µm sterile filters, sediment was transferred to sterile tubes, and sponges were rinsed with filter-sterilized surface seawater and placed in sterile bags. All samples were stored at -20°C on board, then transferred to -80°C storage at the University of Galway and later at the University of Plymouth on dry ice.

**Table 1.**
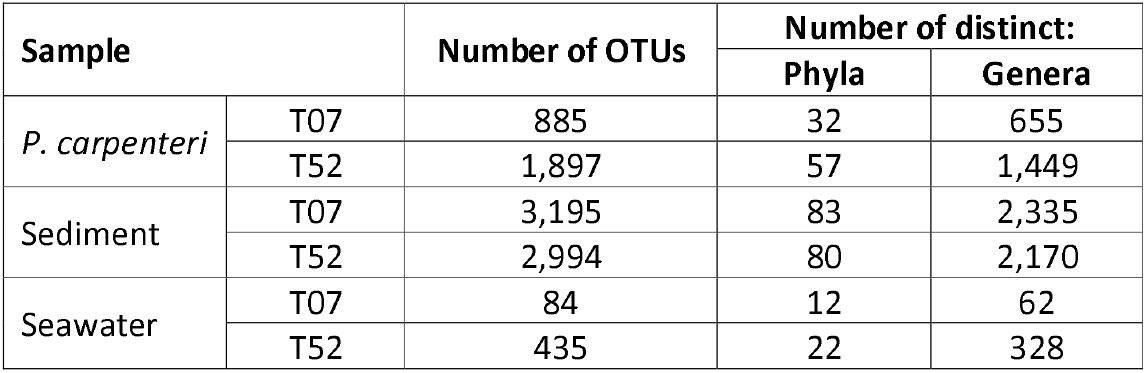
Summary of the number of OTUs generated at phyla and genus level.

| Sample |  | Number of OTUs | Number of distinct: |  |
| --- | --- | --- | --- | --- |
|  |  |  | Phyla | Genera |
| *P. carpenteri* | T07 | 885 | 32 | 655 |
|  | T52 | 1,897 | 57 | 1,449 |
| Sediment | T07 | 3,195 | 83 | 2,335 |
|  | T52 | 2,994 | 80 | 2,170 |
| Seawater | T07 | 84 | 12 | 62 |
|  | T52 | 435 | 22 | 328 |

**Figure 1.**
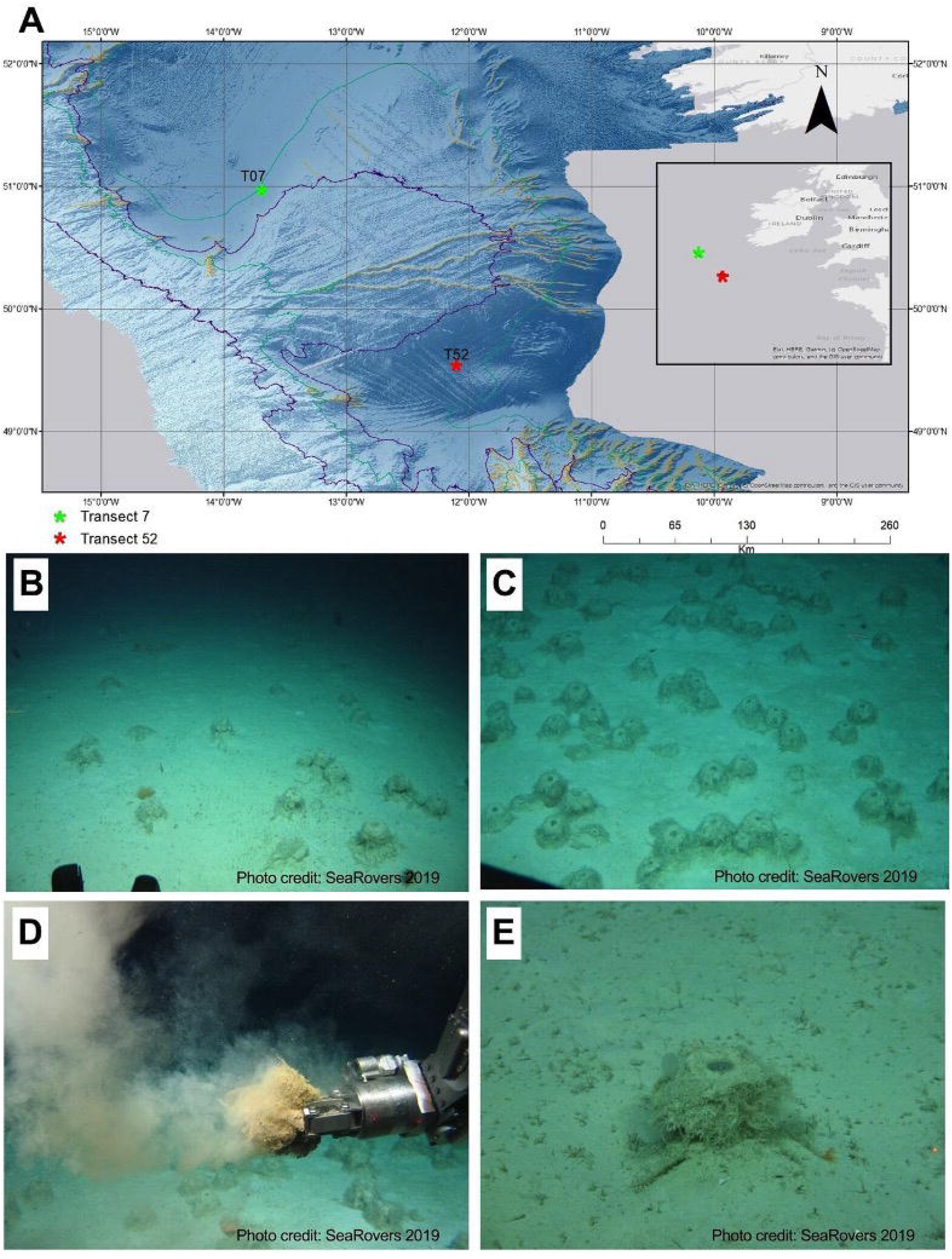
Sampling from *Pheronema* sponge grounds. (A) Map of locations in the North-East Atlantic where *P. carpenteri* sponge samples were recovered for microbiome analysis. The map was generated using ArcGIS. Pheronema sponge grounds are shown from site (B) T07 and (C) T52. (D) Pheronema sponges were collected by ROV. (E) An example of a single *P. carpenter* sponge. Photos are credited to the Marine Institute of Ireland, Galway and were taken on the SeaRovers 2019 cruise CE19-15-029. **Alt Text:** A map showing locations of the sampling sites used in this study and photographs of sponge aggregations and an individual sponge.

### Metagenomic DNA extraction

With a sterile scalpel under a laminar flow hood, the sponge’s outer dermal layer was removed, and mesohyl tissue was cut into 0.25 and 0.5 g portions. DNA was extracted using the DNeasy® Powersoil® Kit (Qiagen) following the manufacturer’s instructions. Extractions with 0.25 g of tissue yielded higher dsDNA concentrations than 0.5 g, which appeared to overload the bead-beating tubes. All extractions were performed in triplicate (n = 3–9) and pooled to compensate for low yields. Water filter samples were quartered and processed individually before pooling, while 0.25 g of sediment was extracted using the same protocol.

DNA purity and concentration were initially assessed using a NanoDrop™ 2000 spectrophotometer (ThermoFisher Scientific) and quantified more accurately with a Qubit dsDNA assay (ThermoFisher Scientific). Fragment lengths and suitability for Oxford Nanopore (ONT) long-read sequencing were evaluated using Agilent Bioanalyzer High Sensitivity DNA chips. Samples with a median fragment length ≥3 kbp were selected for library preparation and sequencing.

### 16S rRNA gene amplicon library and sequencing

The ONT 16S Barcoding Kit (SQK-RAB204) was used following the manufacturer’s instructions, with PCR cycle modification. Ten nanograms of HMW dsDNA from *P. carpenteri* were amplified using barcoded 16S rRNA primers (27F and 1492R) (Heuer *et al*. 1997; Frank *et al*. 2008), on a Veriti 96-Well Thermal Cycler (Applied Biosystems, UK). PCR conditions were: initial denaturation at 95°C for 1 min; 30 cycles of 95°C for 20 s, 55°C for 30 s, 65°C for 2 min; and a final extension at 65°C for 5 min. The number of cycles was increased from 25 to 30 to improve amplicon yield given low dsDNA concentrations and expected low bacterial abundance. Clean-up was performed using AMPure XP beads (Beckman Coulter, UK) and freshly prepared 70% ethanol. Library 1 (L1) was confirmed on an Agilent Bioanalyzer to have a fragment size of ∼1,400 bp. Sequencing was conducted on a MinION™ MkIb using R9 flowcells (FLOWMIN-106), with a 75 µL library loaded through the SpotON port. A standard 48 h protocol was run in MinKNOW on a Dell desktop (32 GB RAM, Intel® Core™ i5-7500 CPU, 3.40 GHz) using the high-accuracy basecalling model.

### Bacterial community composition based on 16S rRNA gene amplicon sequencing data

Bioinformatics analysis was conducted using the Cloud Infrastructure for Microbial Bioinformatics (Connor *et al*. 2016). Demultiplexing was performed by using Porechop v0.2.3 (Wick and Volkening 2018), and reads filtered to a minimum quality (Q>8) by NanoFilt v2.6.0 (Coster *et al*. 2018). Barcodes and adaptors were trimmed using Porechop. After trimming, reads were filtered and cropped to 1,400 bp using Trimmomatic v0.39 (Bolger, Lohse and Usadel 2014). Reads were dereplicated and chimeras detected using VSEARCH v2.14 (Rognes *et al*. 2016), and classified by Kraken2 v2.0.7-beta using the SILVA 16S rRNA v138 database (Wood and Salzberg 2014; Yilmaz *et al*. 2014; Wood, Lu and Langmead 2019) (Wood and Salzberg 2014; Yilmaz *et al*. 2014; Wood, Lu and Langmead 2019). Kraken2 classifications were exported with R package Pavian, with general summary tables of Operational Taxonomic Unit (OTU) counts and taxonomy. Taxonomy and representative sequence files were imported into R studio and used as an input for the package *phyloseq* (McMurdie and Holmes 2013). Phyloseq was used for beta and alpha diversity analysis, and generating community composition figures.

### *P*. *carpenteri* sponge metagenome library

Due to low metagenomic DNA yields, sponge DNA was pre-amplified using the GenomiPhi™ V3 DNA Amplification Kit (GE Healthcare, 25-6601-24) following the manufacturer’s instructions. Initial amplification increased DNA concentration from 4.7 to 6.1 ng/µL; extending the reaction time at 30°C from 1.5 to 2 h increased yield to 28 ng/µL. The ONT Ligation Sequencing Kit (SQK-LSK110) was used to prepare the metagenomic library. DNA was end-repaired and dA-tailed using the NEBNext® Companion Module (cat # E7180S), sequencing adaptors were ligated, and the library was cleaned and assembled. The final library consisted of 250 ng DNA, 4.5 µL nuclease-free water, 25.5 µL loading beads, and 34 µL sequencing buffer. Libraries were sequenced for up to 48 hours with high-accuracy basecalling.

### Bacterial community composition and functional gene analysis of metagenomic sequencing data

Raw ONT metagenomic reads were assembled with metaFlye (Kolmogorov *et al*. 2020), retaining contigs greater than 500 bp that were used for subsequent analysis. Quality of contigs was assessed using metaQuast v2.2 (Gurevich *et al*. 2013; Mikheenko, Saveliev and Gurevich 2016). Open reading frames (ORFs) were predicted using Prodigal v2.6.3 with the metagenomic parameter (*-p meta*) (Hyatt *et al*. 2010). Taxonomic classification of contigs was predicted by Kraken2 using a pre-built database of RefSeq complete or representative genomes. Both taxonomically classified and unclassified contigs were used for further analysis.

Putative genes from BGCs were identified and compiled through comprehensive BLASTp searches (E-value ≥ 1x10^-5^) using DIAMOND v. 2.0.14 (Buchfink, Xie and Huson 2014) against the MiBIG database (Buchfink, Xie and Huson 2015; Kautsar *et al*. 2020). A reciprocal BLASTp search of identified sequences against a database of Sponge 29 metagenomic proteins and MiBIG database was carried out to identify Reciprocal Best Hits (RBH). RBH were used to classify sequences before phylogenetic analysis. Functional annotation of key enzymes was conducted by annotating ORFs with four curated databases using DIAMOND for a BLASTp algorithm. Databases included: NaPDoS (Natural Product Domain Seeker; accessed Oct 2020) (Ziemert *et al*. 2012), NCycDB (nitrogen metabolic cycling pathways; accessed Jan 2021) (Tu *et al*. 2019), UniProtKB (UniProt Knowledgebase; accessed Jan 2021) (Bateman and Consortium 2018), antiSMASH v5 (secondary metabolite analysis; accessed Jan 2021) (Kautsar *et al*. 2020). Homologous sequences were identified when the BLASTp search fulfilled the minimum parameters of E-value ≤ 10^-5^, and a bit-score > 50 (Pearson 2013).

### Statistical Analysis

Two-Way ANOVA and pairwise comparisons of the relative abundance of taxa between sampling sites were performed using GraphPad Prism v. 9.0.1 for Windows (GraphPad Software, San Diego, California USA, www.graphpad.com). Graphical outputs were generated using either GraphPad Prism or R-studio.

## Results

### DNA extraction, library preparation, sequencing, and raw read processing

Extracting sufficient DNA from *P. carpenteri* mesohyl tissue was challenging, yielding on average 1,072.5 ng dsDNA per g of wet tissue. Bioanalyser fragment analysis showed most samples contained 3–6 Kbp fragments, suitable for 16S rRNA metabarcoding but suboptimal for metagenome sequencing. Water-filter samples were highly fragmented and had the lowest DNA yields. Nine sponge samples with sufficient DNA were pooled into Library 1 (L1). Samples with low DNA yields required additional amplification and pooling into Libraries 2 and 3, achieving a maximum of 13 ng dsDNA per sample for sequencing. Sequencing generated ∼5.6 million reads, of which 2.4 million (42%) passed quality, length (>1,400 bp), and chimera filters. Kraken2 classification identified 3,571 OTUs across all samples, spanning 85 phyla and 2,626 genera (Table 2). Sediment samples had the highest OTU richness, followed by *P. carpenteri*, then seawater. Rarefaction analyses indicated adequate sampling depth for sediment and T52 sponge samples, while seawater and T07 sponge samples were under-sampled (Supp. Figure 1). Overall, T52 sponges exhibited higher OTU diversity and read abundance than T07.

### Spatial Variation in Prokaryotic Communities of *Pheronema* Aggregations and Surrounding Sediments

The *P. carpenteri* bacterial community is represented by a high abundance in Alpha-proteobacteria, Gamma-proteobacteria, Actinobacteria, and Planctomycetes. Members of the rare phyla (below 0.1% relative abundance) included, Firmicutes, Nitrospirota, Gemmatimodoat, and Spirochaeota (Figure 2). Greater overall diversity was seen in the sediment than the sponges, with sediment samples being relatively uniform between the two transect sites concerning diversity, evenness and abundance of reads (Supp. Figure 1B-D). This was not the case for sponge samples, where the two *Pheronema* aggregations were distinct. Sponges from the aggregation at T52 generated the greater mean community evenness and diversity (Supp. Figure 1D).

**Figure 2.**
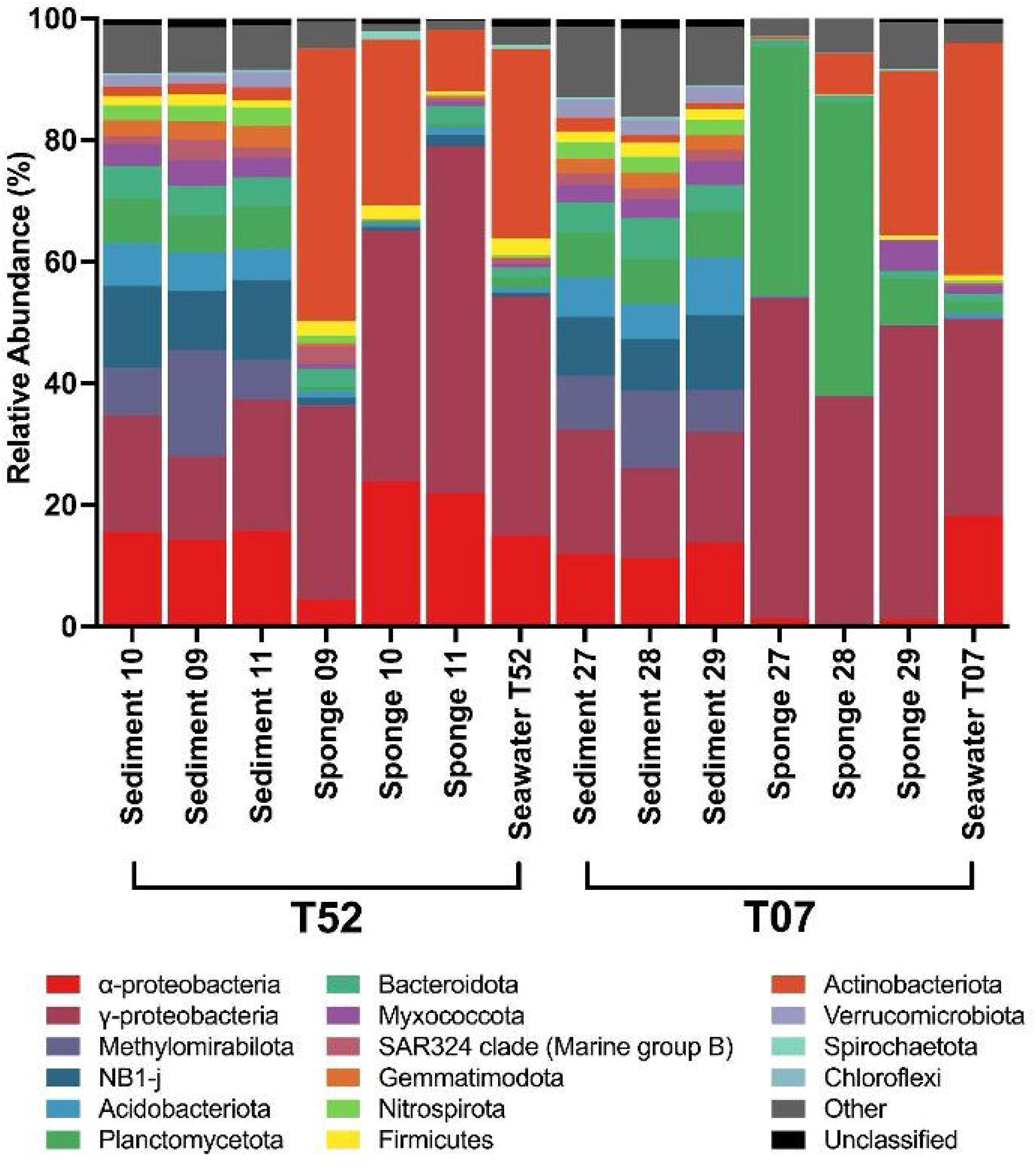
The relative proportions of the most abundant bacterial phyla in *P. carpenteri* aggregations, sediment, and water samples from sites T07 and T52 based on 16S rRNA gene amplicon profiling. Only the most abundant taxa are shown, with phyla below 0.1% among the samples compiled into ‘Other.’ Proteobacteria is split into classes (α, Alphaproteobacteria; and γ, Gammaproteobacteria). **Alt Text:** A stacked bar chart showing the abundance of bacterial phyla revealed by sequencing 16S rRNA gene amplicons that were generated from DNA isolated from sponge samples

Proteobacteria, Methylomirobilota, members of the NB1-j group, and Acidobacteriota were the most abundant phyla in sponge individuals. Seawater samples had a higher abundance of Proteobacteria and Actinobacteria, and shared a greater resemblance to sponge samples than the sediment. The differences between the bacterial compositions in sponge individuals was explored (Figure 3). Major differences between aggregations were observed in the mean relative abundance of Alphaproteobacteria (T07, mean = 0.948 ± 0.67%; and T57, mean = 16.71 ± 10.7%; p = 0.038), Actinobacteria (T07, mean = 11.37 ± 13.97%; and T57, mean = 27.45 ± 17.36%; p = 0.032), and Planctomycetes (T07, mean = 32.29 ± 21.68%, and T57 mean = 0.53 ± 0.15%; p <0.0001), as calculated by a two-way ANOVA (Figure 3B; Supp. Table 3). Sediment samples showed no significant differences between aggregation sites based on beta diversity analyses (Supp. Figure 2; Supp. Table 3), with the exception of the collective phylum category ‘Other,’ which differed between T07 (mean = 11.59 ± 2.50%) and T52 (mean = 7.53 ± 0.35%; p = 0.029, Figure 3A-B). The same comparisons could not be conducted with the seawater samples due to a lack of sequencing replicas, as a result of pooling water samples together.

**Figure 3.**
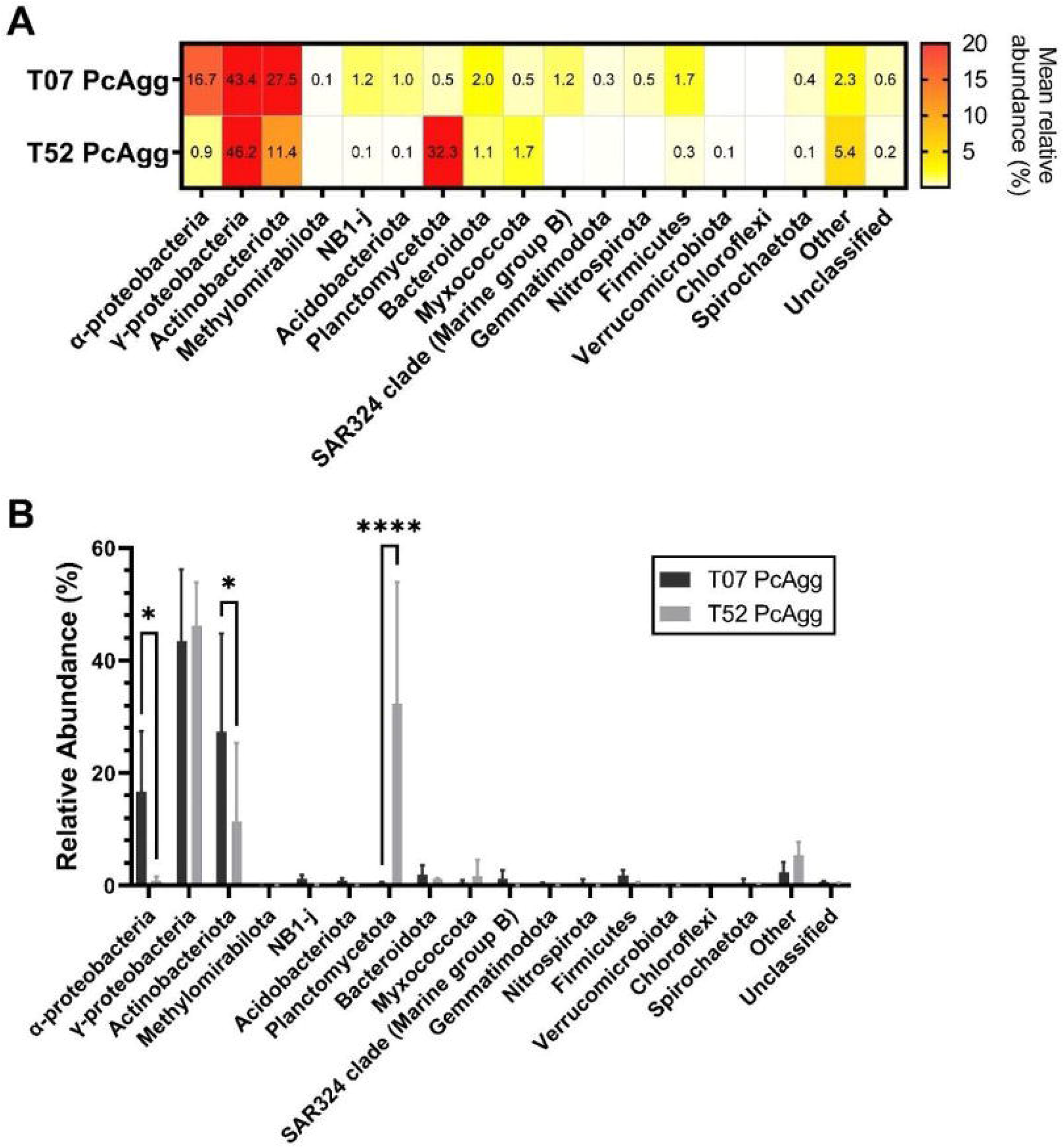
Detailed comparison of the prokaryotic composition shows Planctomycetes, Actinobacteria, and Alphaproteobacteria relative abundances are significantly different between sponges from transect sites T07 and T52 *P. carpenteri* aggregations (PcAgg). (A) Mean relative abundance at phyla level, mean relative abundances > 0.1% are shown. (B) Comparing the relative abundances between two *P. carpenteri*, data are shown as mean ± SD (N = 3). (Significant p values from a pairwise comparison are shown in figure: ⍰1, p < 0.05; ⍰, p < 0.0001). **Alt Text:** A heatmap and bar chart showing relative abundances of bacteria in sponge samples from two sampling sites in the North Atlantic

In addition to differences observed between *P. carpenter* from different sites, there were notable differences between the biological replicates from the same *P. carpenter* sample. This was noted when comparing the relative read abundances between samples of the same *P. carpenteri* (Figure 3). Ternary plots were used to examine bacterial phyla distribution between the *P. carpenteri* individuals of the same sampling site (Figure 4A). As indicated, specific taxa were nearly exclusive to a single sponge individual from each site, such as Myxococcota and Spirochaete, for Sponge 10 (T52) and Sponge 29 (T07), respectively. Proteobacteria were evenly distributed at T07 but less so at T52. Patescibacteria, Actinobacteria, and Bacteroidetes were unevenly distributed at both sampling sites.

**Figure 4.**
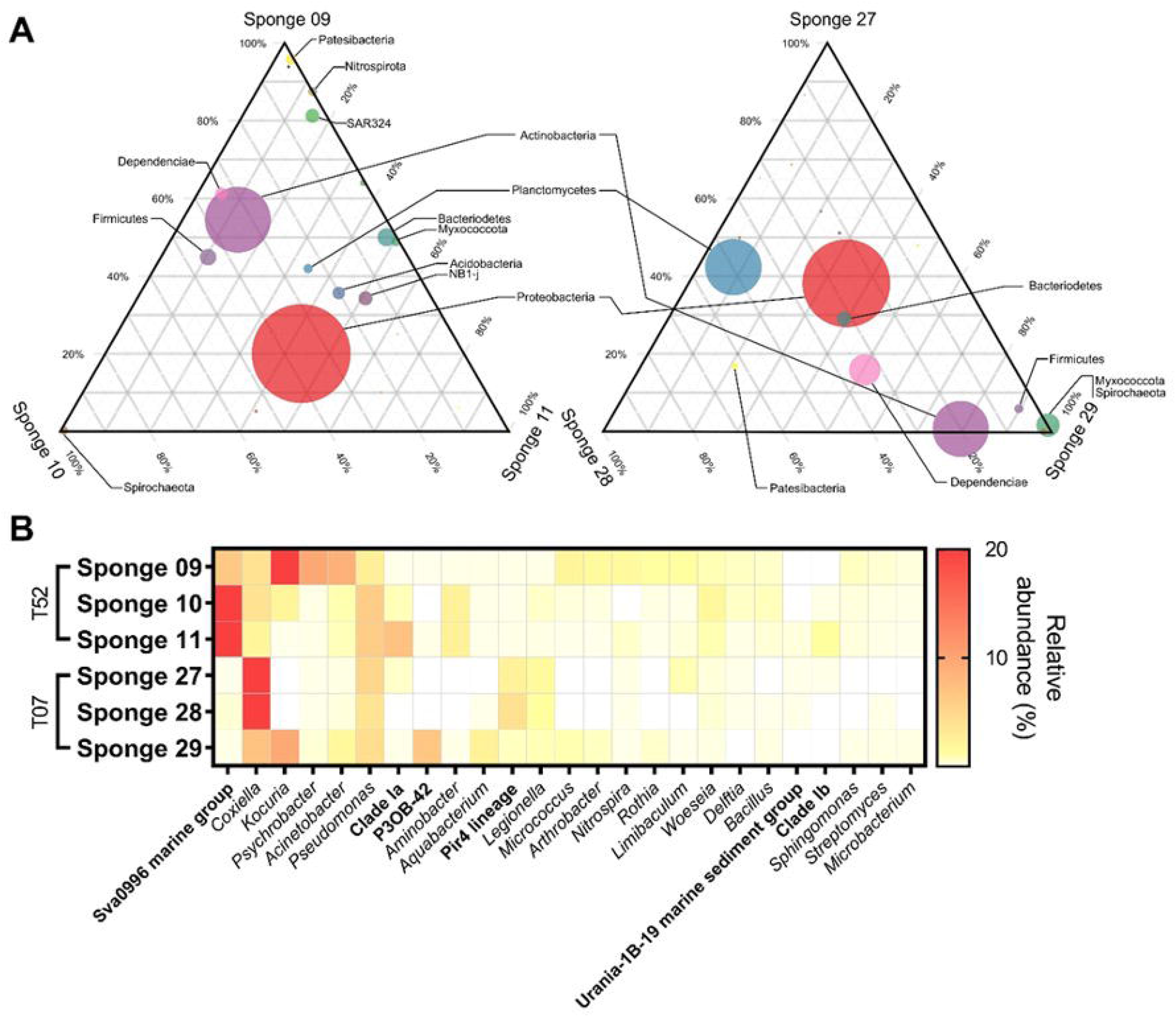
Distribution of bacterial taxa between sponge individuals. (A) Ternary plot of the most abundant bacterial phyla. The left ternary plot is for T52 samples, and the right is T07 samples, and circle size indicates the relative abundance of each phylum. (B) Heatmap of relative abundance of genus-level taxa between sponge samples. **Alt Text:** Ternary plots and a heatmap showing the variation in bacterial populations between different replicate samples of the same sponges recovered from the North Atlantic

At the genus level, *Coxiella, Kocuria, Pseudomonas, Woeseia*, and Pir4 lineage were present in all individuals. T52 sponges were rich in Sva0996 marine group bacteria, which were present at lower relative abundances in T07 *P. carpenteri* sponges (Figure 4B). *Aquabacterium* and Pir4 lineage abundances were uniform for T07 sponges but not for T52. Lack of uniformity between biological replicas of the same aggregation can be seen in various taxa. Greater overall differences in the taxa were observed from T07 sponges as compared to T52 sponges, which were far more uniform. *Aquabacterium, Arthrobacteri*, Clade Ib, *Kocuria, Microbacterium, Micrococcus, Rothia* and *Sphingomonas* were all present in a single sponge sample from T07. In contrast, the Urania-1B-19 marine sediment group was the only taxon unique to a single sponge sample (Sponge 11) from T52. More significant dissimilarity was seen between sediment samples and the seawater and sponge samples measured by Bray-Curtis Dissimilarity clustering (Figure 5C). UniFrac weighted component analysis shows a clustering of Sponge T52 individuals with both seawater samples (Figure 5B) while also showing a tight clustering of sediment samples from T07 and T52.

**Figure 5.**
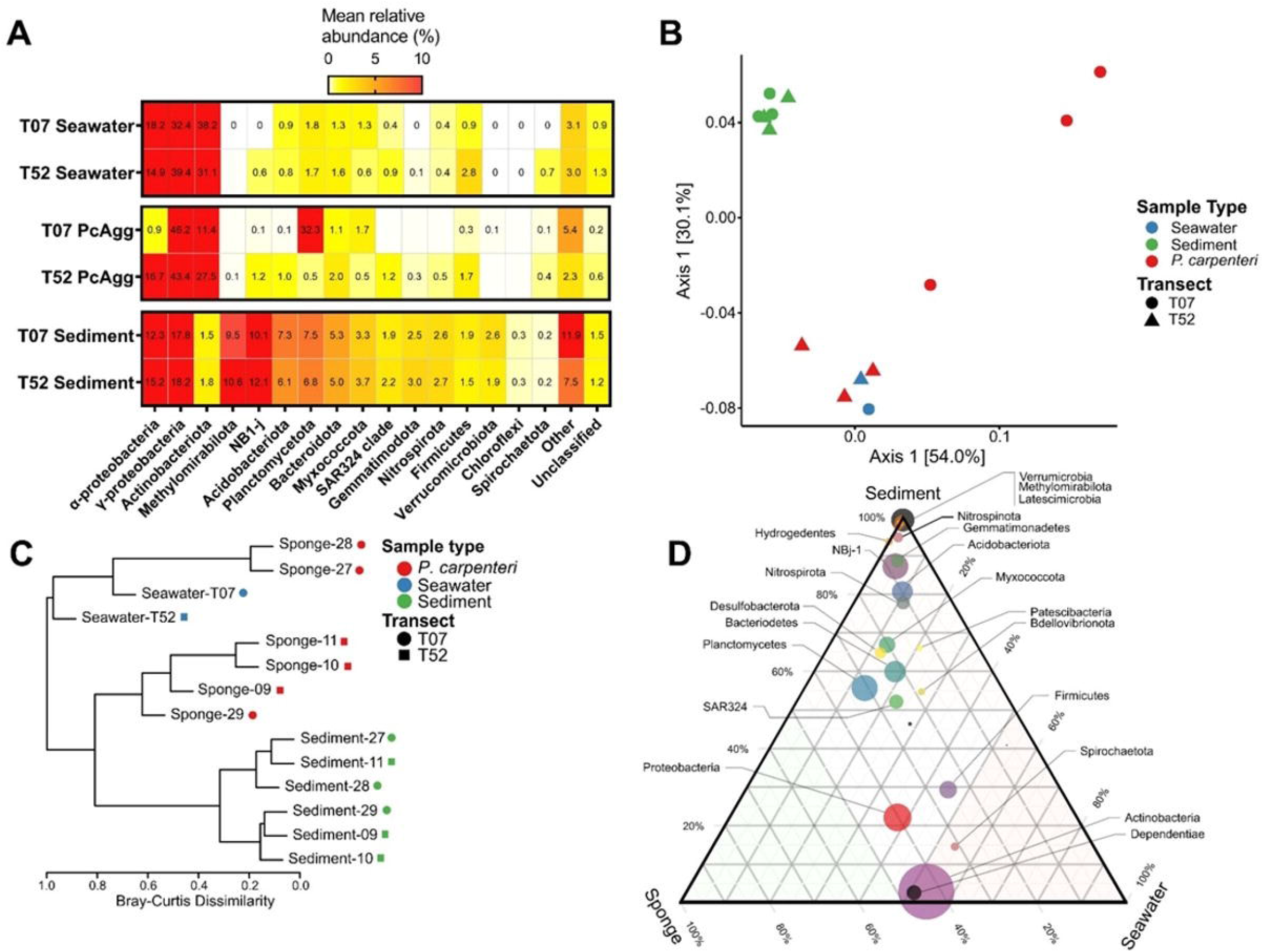
Sponge and seawater samples show closer resemblance, while *P. carpenteri* (PcAgg) sampling site and individual taxa are significant factors shaping the bacterial community composition of *P. carpenteri*. Comparison between the relative abundances of (A) all samples at the phylum level and (D) only sponge samples at the genus level. (B) Component analysis using weighted uniFrac distances. (C) Bray-Curtis dissimilarity dendrogram. (D) Ternary plot of pooled samples by biotypes of the most abundant phyla. **Alt Text:** Various plots and a phylogenetic tree used to describe the relationships between bacterial groups identified in samples of sponge, sediment and seawater from two sites in the North Atlantic.

Ternary plots were used to examine bacterial taxa distribution between the pooled *P. carpenteri*, sediment, and seawater samples (Figure 5D). As shown in the ternary plot, Proteobacteria, Actinobacteria, Dependentiae, Firmicutes, and Spirochaetota, which make up the most abundant taxa in the *P. carpenteri* sponge samples, were evenly distributed between sponge and seawater. The remaining taxa were far more abundant in sediment samples. Planctomycetes and SAR324 Marine Clade were evenly distributed between all sample types (i.e. sediment, sponge and water). Taxa rarely observed in the sponges were more common in sediment, such as Nitrospirota, Nitrospinota, Patescibacteria.

### Potential functions of microbial members of *P*. *carpenteri*

Sponge 29 was chosen to sequence the metagenome because it had a high enough dsDNA concentration to meet the Ligation Sequencing Kit’s required starting material (1 µg of dsDNA). Sponge 29 was sequenced on a 48 h sequence cycle and, using a whole flow cell, generated 232,000 reads. These reads were then cleaned (minimum lengths > 150 bp), from which 3,222 contigs were assembled using metaFlye (min 501 bp, max 49 Kbp). The NCBI Sequence Taxonomic Analysis Tool (STAT) classified 3.64% of the reads as bacterial, while 96.36% remained unclassified. Because reference genomes are not available for *P. carpenteri* or its closest relatives, we are unable to determine what proportions of the unclassified reads derive from sponge genomic material or from dark microbial matter.

From the 3,227 contigs, 24,260 protein-coding regions were predicted by Prodigal. Translated amino acid sequences were parsed using BLASTp against both the NaPDoS and NCyc database. There were 21 Best Reciprocal Hits (BRH) against the NaPDoS database (Supp. Table 3). Against the NaPDoS, the vast majority of BRH were for Fatty Acid Synthesis (FAS) and some Type II PKS. Besides fatty acids, additional predicted products for some of these pipelines were Aclacinomycin, Alnumycin, and spore pigments. When the contigs were parsed through the antiSMASH v.6 online tools, four clusters were identified: a Type I Polyketide Synthase (T1PKS) region, a Ribosomally synthesised, and post-translationally modified peptide-like (RiPP-like) region, and two Arylpolyene regions (Supp. Figure 4). Contig_3500 showed low similarity (15%) to the NRP+T1PKS cluster producing nosperin and related to pederin group members (Kampa *et al*. 2013). Pederin group members have only been identified from non-photosynthetic bacteria associated with marine sponges and beetles (Robinson *et al*. 2007).

More success was found when parsing the predicted proteins dataset against the NCyc database with 375 BRH identified homologues (Supp. Figure 5, Supp. Table 4). The majority of BRH were proteins classified under the denitrification pathway, including 110 BRH homologues to Nitrite reductase (NO-forming, *nirK*). Most of these were identified on contigs which could not be taxonomically classified or were of Proteobacterial origin. A further 64 BRH were Nitrous-oxide reductases (nosZ), and 28 BRH were homologues of Nitrite reductases (nirS). Two nitrogen fixation genes were identified; Nitrogenase iron protein NifH (7 BRH) and Nitrogenase-stabilizing/protective protein NifH (6 BRH). Although this approach will not have fully separated microbial from host-derived genes, there is growing evidence in the literature supporting the involvement of ammonia-oxidizing archaea (AOA) in hexactinellids’ closed-loop nitrogen cycling (Tian *et al*. 2016; Garritano *et al*. 2023, 2024).

## Discussion

Sponges are key players in marine ecosystems, providing essential functions such as nutrient cycling and habitat formation, and supporting biodiversity, with many of these activities being mediated by complex associations with microbiota (Bell 2008; Goeij *et al*. 2013; Maldonado 2016). While the microbiomes of Demospongiae have been extensively studied, Hexactinellida (glass sponges), which inhabit almost exclusively deep-water environments, remain poorly characterized. In the North East Atlantic, *P. carpenteri* forms dense aggregations at ∼1,200 m depth, with spicule mats that support diverse macrofauna and are recognized as Vulnerable Marine Ecosystems under multiple policy instruments (Bett and Rice 1992; Howell *et al*. 2016; Vieira *et al*. 2020).

16S rRNA gene profiling revealed that *P. carpenteri* harbours a relatively low-diversity bacterial community but contains taxa characteristic of both low microbial abundance (LMA) and high microbial abundance (HMA) sponges. As in other Hexactinellida (Steinert *et al*. 2020), Proteobacteria, particularly Alphaproteobacteria, dominated the microbiome, accompanied by Bacteroidetes and Planctomycetes, indicative of LMA traits. Notably, Actinobacteria, typically associated with HMA Demosponges, were enriched in sponges from site T07, suggesting that Hexactinellida may host unique microbial assemblages that complicate the HMA/LMA dichotomy. In *P. carpenteri*, Chloroflexi were present at low abundance, consistent with previous observations in LMA sponges (Moreno-Pino *et al*. 2024).

Significant differences in microbial composition were observed between sampling sites, with Planctomycetes enriched at T52 and Actinobacteria and Alphaproteobacteria enriched at T07 (Figure 5B). This aligns with previous studies showing species-specific and site-specific microbiomes in sponges (Hentschel *et al*. 2003; Thomas *et al*. 2016, 2016; Steinert *et al*. 2020; Busch *et al*. 2022). While trawling can influence sponge microbiomes (Busch *et al*. 2020), no trawling activity was reported at our sampling sites (O’Sullivan, Healy and Leahy 2019), suggesting other environmental or stochastic factors may drive observed differences.

Hexactinellid sponges exhibit greater variation in microbial composition between biological replicates than Demosponges (Steinert *et al*. 2020), which we also observed at both the phylum and genus levels. In our study, we also observed intra-aggregation differences, with sponges from the same aggregation exhibiting taxonomic variation. Differences in OTU counts between T07 and T52 may partly reflect sequencing depth, but also highlight the inherent heterogeneity of hexactinellid microbiomes. This trend has been noted in a much larger study of deep-sea sponge microbiomes, with the authors concluding that deep-sea sponge individuals carry their own set of microbes (Busch *et al*. 2022), as well as species-specific relationships (Garritano *et al*. 2023). Despite these differences, individuals within the same aggregation were generally more similar to one another than to sponges from different aggregations (as observed in comparisons between transect sites T57 and T7), even though the surrounding seawater samples were largely uniform (Figure 5B). These patterns highlight the need to further investigate the relative contributions of aggregation membership and genetic lineages within aggregations to microbial community composition.

Interestingly, *P. carpenteri* microbiomes showed greater similarity to seawater than to sediment samples (Figure 2, Figure 4), consistent with patterns reported in other Hexactinellida (Steinert *et al*. 2020). Despite being embedded in the seabed and coated in sediment, the overlap with sediment microbiota was minimal, suggesting that sediment has little influence on the sponge’s core microbial community. Future comparisons between inner mesohyl and outer dermal layers may provide further insight into tissue-specific microbial distributions, since the inner mesohyl was utilised in this study. Low DNA yields and challenges in 16S rRNA amplicon generation were consistent with the low microbial abundance typical of Hexactinellida, as well as the sponge’s high spicule content.

Metagenomic sequencing of *P. carpenteri* was also limited by low DNA yields, likely due to the small proportion of microbial biomass relative to the extensive siliceous spicule matrix of Hexactinellida and the low microbial abundance characteristic of this species. As a result, sequencing depth was insufficient to robustly identify biosynthetic gene clusters (BGCs) of interest for natural product (NP) discovery. Although a number of PKS and NRPS homologues were detected, the dataset was not comprehensive enough to enable extensive mining of novel compounds.

These challenges highlight that metagenomic approaches alone may be insufficient for characterisation of deep-sea glass sponges like *P. carpenteri*, and that complementary culture-based strategies targeting sponge-associated bacteria may be more practical for NP discovery. Previous work has demonstrated that *P. carpenteri* harbours cultivable bacteria capable of producing bioactive compounds (Koch *et al*. 2021; Hesketh-Best *et al*. 2023; Conway *et al*. 2025), suggesting that focused cultivation combined with genomic screening may provide a more feasible path to accessing the chemical potential of this species. Bacterial species cultivated from *P. carpenteri* include *Micrococcus* spp., *Delftia* spp., and *Psychrobacter* spp., and here are identified in the 16S rRNA amplicon survey (Figure 4B). Interestingly, excepting *Psychrobacter* spp., these genera represented very low abundance taxa in the amplicon survey (Figure 4).

The metagenomic analysis of Sponge 29 revealed a relatively low proportion of reads confidently classified as bacterial, highlighting the persistent challenge of resolving host versus symbiont genomic material in deep-sea sponges where reference genomes remain scarce. The high abundance of *nirK*, alongside *nosZ* and *nirS* homologues, suggests that denitrifying bacteria, primarily associated with Proteobacteria or taxonomically unresolved contigs, may contribute to regulating nitrogen turnover within the *P. carpenteri* holobiont. The detection of *nifH* homologues further indicates the potential for biological nitrogen fixation, implying that both nitrogen loss and gain pathways may coexist in the associated microbiome, as has been observed in hexactinallids *Schaudinnia rosea* and *Vazella pourtalesii* (Maldonado *et al*. 2021).

*P. carpenteri* is an ecologically important deep-sea sponge whose microbiome exhibits features of both LMA and HMA sponges. In this small-scale study, we observed variation in microbial composition between aggregations and individuals, with clear site-specific patterns. The microbiome shares more taxa with seawater than sediment, consistent with other Hexactinellida. Metagenomic analysis was constrained by low DNA yields and limited sequencing depth, restricting the detection of BGCs for natural product discovery. These methodological limitations suggest that culture-based approaches may currently be the most effective strategy for exploring the biotechnological potential of *P. carpenteri*. Given the small sample size and scope of this study, further research is clearly warranted. Expanded sampling across multiple aggregations, coupled with metatranscriptomic analyses and targeted assessment of archaeal communities, will be essential to fully resolve the ecological functions and biotechnological potential of this important deep-sea species.

## Data availability

All quality controlled 16S rRNA amplicons and the Nanopore metagenome reads are available through the Sequence Reads Archive (SRA) repository under the BioProject accession PRJNA885129, individual SRA accessions can be found in Supplemental Table 1.

## Author Contributions

Study conceptualised, devised, planned, conducted by all authors. Sponges were collected by KLH. Data curation, analysis and visualizations were carried out by PJHB and MJK. Writing of the original draft was performed by PJHB and MJK, and all authors contributed to the reviewing and editing of the manuscript. Supervision and funding acquisition was by MU, KLH, PJW and NF.

