## Supplementary material for "A metagenomic exploration of the bacterial community composition of two deep-sea *Pheronema carpenteri* sponge aggregations from the North Atlantic; insights into ecosystem services": Supp. Figure

**Supplementary Table 1 –** Quality and quantity of metagenomic DNA samples. Molarity and media fragment size determined by the BioAnalzyer, concentration assessed by Qubit.

**Supplementary Table 2 –** Summary of Two-Way ANOVA multiple comparison test for data presented in Supplementary Figure 4B.

**Supplementary Table 3 –** BLASTp tabular output for the Best Reciprocal Hits (BRH) against NaPDoS database. Diamond BLASTp of predicted proteins from metagenomic data against the NaPDos database. (pident, Percentage of identical matches (%); length, alignment length; mismatch, Number of mismatches; gapopen, Number of gap openings; qstart, Start of alignment in query; qend, End of alignment in query; sstart, Start of alignment in subject; send, End of alignment in subject; evalue; Expected value; bitscore, Bit score).

**Supplementary Table 4 –** Extended BLASTp tabular output for the Best Reciprocal Hits (BRH) against NCyc database. Diamond BLASTp of predicted proteins from metagenomic data against the NaPDos database. (pident, Percentage of identical matches (%); length, alignment length; mismatch, Number of mismatches; gapopen, Number of gap openings; qstart, Start of alignment in query; qend, End of alignment in query; sstart, Start of alignment in subject; send, End of alignment in subject; evalue; Expected value; bitscore, Bit score)

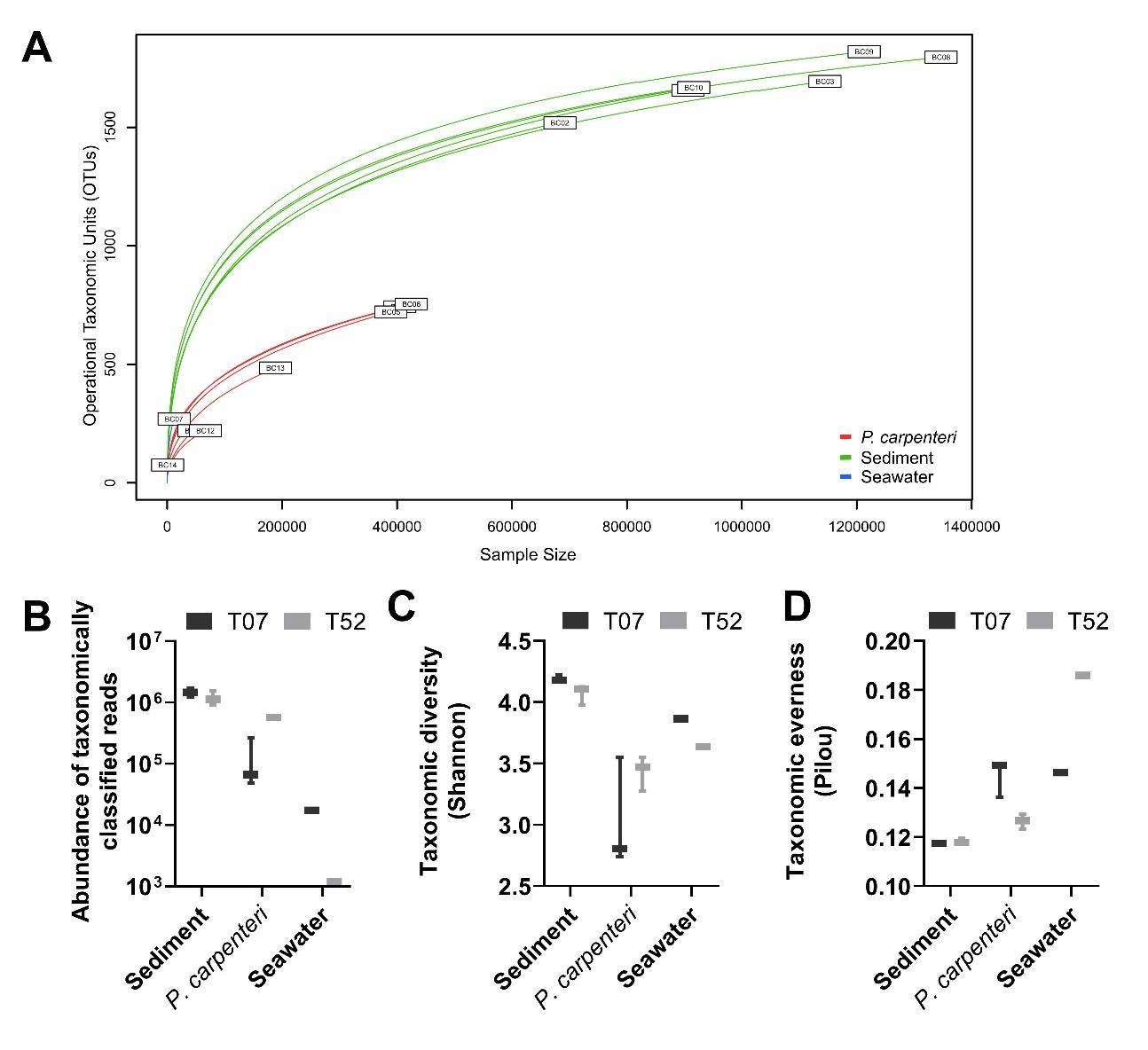

**Supplementary Figure 1 –** *P. carpenteri* exhibit a low observed and Shannon diversity compared to sediment communities, on a par with water column diversity. (A) Rarefraction curve, (B) abundance of taxonomically classified reads, (C) taxonomic diversity as calculated by Shannon diversity metric, (D) taxonomic evenness as calculated by Pilou evenness metric. (A-D) *P. carpenteri*, sediment and water samples are grouped by sampling sites.

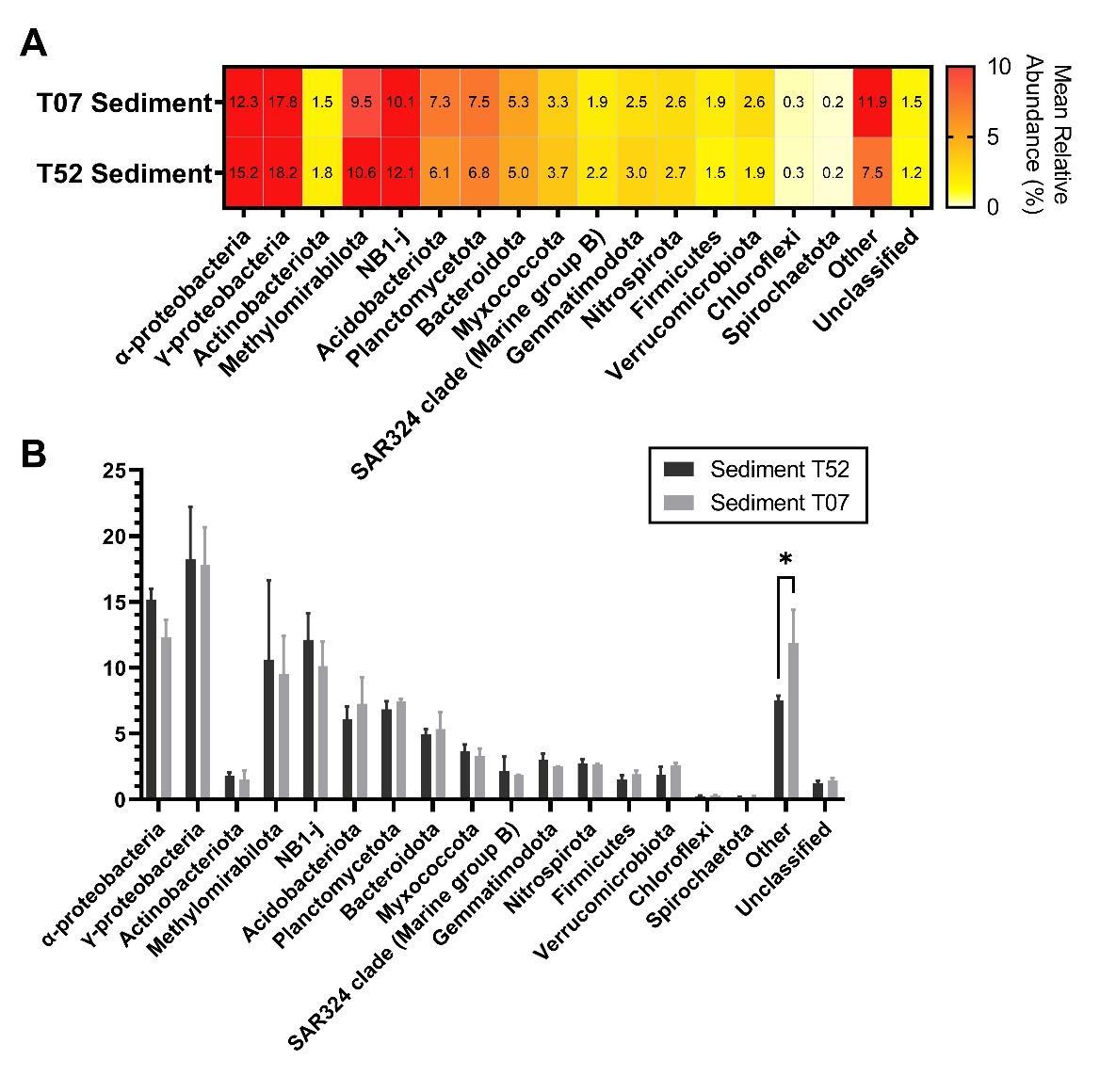

**Supplementary Figure 2 –** A comparison of sediment sampling sites reveals that rare taxa (< 0.1%) differentiate the two sites. (A) Mean relative abundance at phyla level, mean relative abundances > 0.1% are shown. (B) Comparison of the relative abundances between two *P. carpenter* samples, data are shown as mean ± SD (N = 3). Full results of Two-Way ANOVA can be found in Supplemental Data File (Significant *p* values from a pairwise comparison are shown in figure: ∗, p < 0.05).

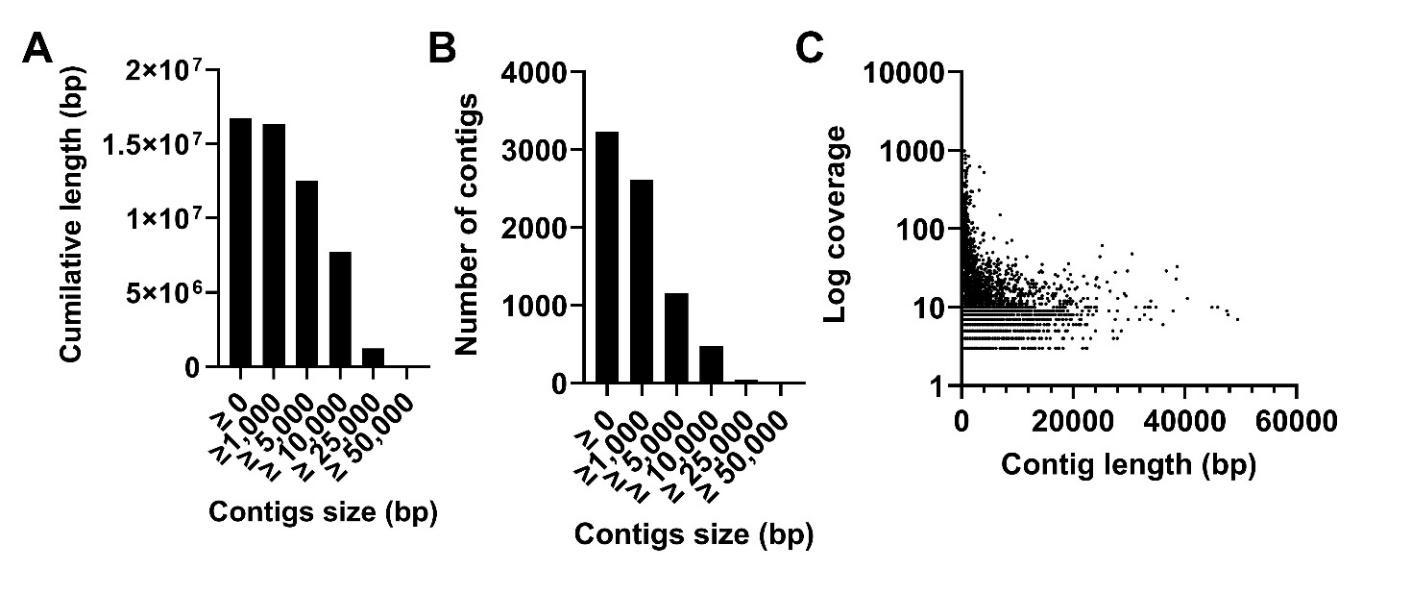

**Supplementary Figure 3 –** Quality analysis of contigs generated from Sponge 29 metagenome using metaFlye. Summary of contigs was generated by using QUAST, (A) the cumulative lengths of contigs binned by contig length, (B) the number of contigs binned by length, and the coverage of contigs by length.

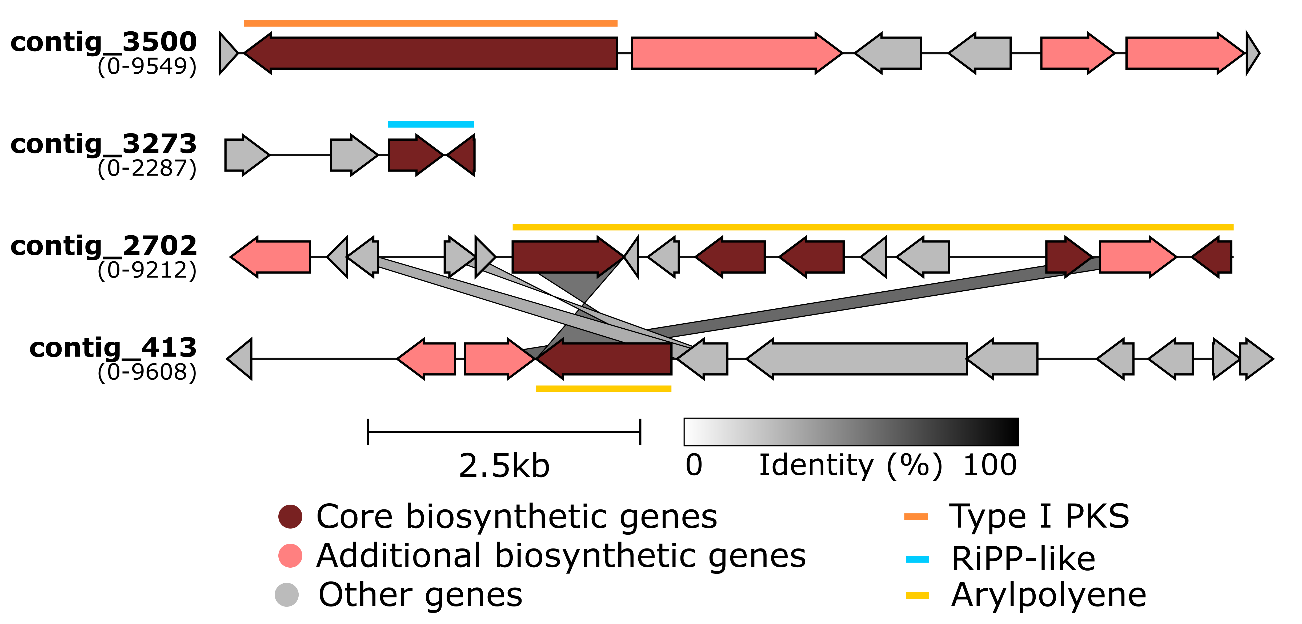

**Supplementary Figure 4 –** antiSMASH v.5 cluster predictions for Sponge 29. antiSMASH results were downloaded as GenBank (.gbk) file and aligned, and cluster arrow map was visualised using Clinker*,* annotations were added using Inkscape. Similar regions between the two Arylpolyene clusters are indicated. The coloured bars above arrows represent the region identified as belonging to a particular class of BGC. The start and end region of the cluster is indicated in the parentheses below the contig name. Genes with similar proposed functions are indicated with identical colours.

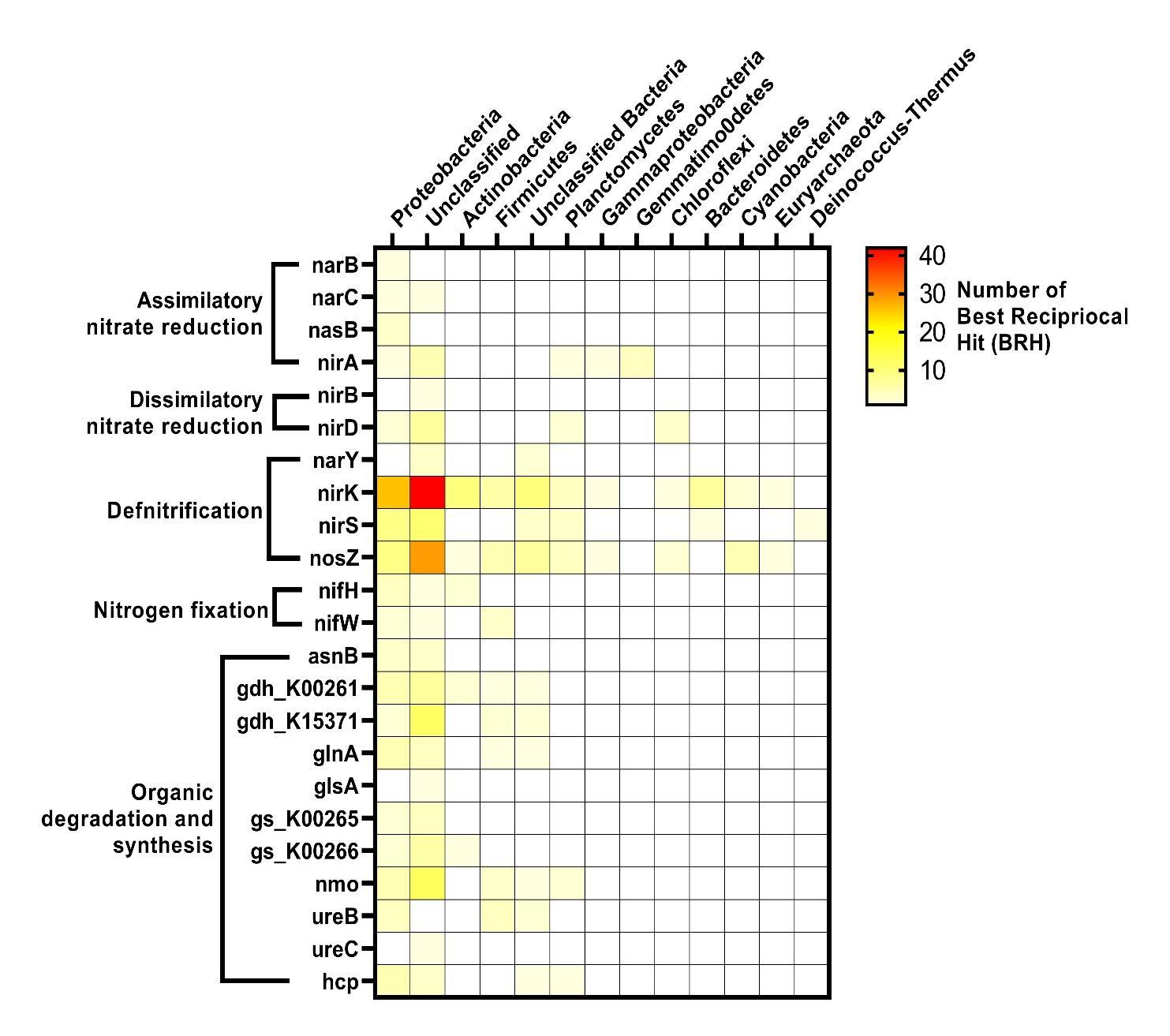

**Supplementary Figure 5** – **Specific functions abundant in *P. carpenteri*-associated bacterial metagenome sequence contigs annotated with COG.** The brightness (red) in the heatmap reflects the abundance (number of BRH) of a particular protein in the Sponge 29 metagenome against the NCyc database, white indicates absence. Functional pathways are indicated to the left of the gene name. Full list of COGs identified can be found in Supplementary Table 4.
